# Temporal and environmental factors interact with rootstock genotype to shape leaf elemental composition in grafted grapevines

**DOI:** 10.1101/2022.02.28.482393

**Authors:** Zachary N Harris, Julia E Pratt, Niyati Bhakta, Emma Frawley, Laura L. Klein, Misha T Kwasniewski, Zoë Migicovsky, Allison J Miller

## Abstract

Plants take up elements through their roots and transport them to their shoot systems for use in numerous biochemical, physiological, and structural functions. Elemental composition of above-ground plant tissues, such as leaves, reflects both above- and below-ground activities of the plant genotype, as well the local environment. Perennial, grafted plants, where the root system of one individual is fused to the shoot system of a genetically distinct individual, offer a powerful experimental system in which to study the role of the root system in the elemental composition of the shoot system. We measured elemental composition of over 7000 leaves in the grapevine cultivar ‘Chambourcin’ growing ungrafted and grafted to three rootstock genotypes. Leaves were collected over multiple years and phenological stages (across the season) and along a developmental time series. Temporal components of this study had the largest effect on leaf elemental composition; and rootstock genotype interacted with year, phenological stage, and leaf age to differentially modulate leaf elemental composition. Further, the local, above-ground environment affected leaf elemental composition, an effect influenced by rootstock genotype. This work highlights the dynamic nature by which root systems interact with shoot systems to respond to temporal and environmental variation.

## Introduction

Through their roots, plants take up elements from the soil and incorporate them throughout the plant body for use in biochemical reactions and physiological and structural functions. Generally, elements are classified into two broad categories: macronutrients, such as phosphorous and magnesium, are needed in high concentrations for nucleic acid structure and photosynthesis (Marschner, 2011; de Bang et al., 2021); while micronutrients, such as nickel and molybdenum, are needed in much smaller amounts as enzyme cofactors used in, for example, amino acid and ammonia metabolism (Hänsch and Mendel, 2009; Marschner, 2011). Plant elemental composition is influenced by genetic and environmental factors, but relationships of climatic variation with temporal shifts in elemental composition, remain undescribed.

The traditional model for plant elemental accumulation posits a dynamic source/sink system where plant roots take up macro- and micronutrients from the soil (a source) during early development and shuttle these elements to young leaves (a sink) via xylem (Batten et al., 1986; Lynch and White, 1992; Marschner, 2011). Over the course of the growing season, young leaves act as a reservoir (a new source) for elements needed elsewhere in the plant such as fruit or seeds (a new sink) via the phloem. Variation in the elemental composition of plant tissue is governed by genetics, the local environment, and their interaction (GxE). Numerous studies have shown that elemental accumulation is under quantitative genetic control within a given environment (Ziegler et al., 2018; Fikas et al., 2019; Liu et al., 2020; Cobb et al., 2021); however, this effect varies across different environments. For example, in seeds of *Sorghum bicolor*, most of the 20 elements assayed showed significant variation explained by the interaction of genotype and environment, but the strength of the GxE effect varied by element (Shakoor et al., 2016). Similarly, in *Zea mays*, 80% of the quantitative loci correlating with elemental composition in kernels were site specific (Asaro et al., 2016). These studies suggest that variation in elemental composition is complex and is controlled by both genetic and environmental regulation.

Current understanding of plant elemental composition is based primarily on annual species that complete their life cycles within one year. Annual species follow a pattern where early season growth and resource allocation take place in vegetative tissue and later season growth and resource allocation target reproductive tissue, then followed by senescence and death (Friedman and Rubin, 2015). In contrast, perennials allocate resources to vegetative and reproductive tissue in a cyclical fashion over the course of multiple years (Friedman and Rubin, 2015; Friedman, 2020). Between cycles, deciduous perennial species drop leaves and experience periods of dormancy in preparation for the next cycle of vegetative growth, a process which requires mechanisms for mineral reallocation. Leaves of perennial species experience a sharp decrease in nitrogen, phosphorus, and potassium prior to leaf drop (Sánchez Alonso and Lachica, 1987a, 1987b); whereas concentrations of calcium and manganese increase in leaves prior to senescence, often attributed to low phloem mobility (Sánchez Alonso and Lachica, 1987a, 1987b; Roca-Pérez et al., 2006). Variation exists in phloem mobility of certain elements, with low mobility proposed for boron in *Chamelaucium* spp. (Maier and Chvyl, 2002) and magnesium in *Vitis vinifera* (Navarro et al., 2008; Holzapfel, 2019). These patterns are reflected in elemental composition in leaf position along a shoot (George et al., 1989; Maier et al., 1995; Huber et al., 2016), generally a correlate for leaf age. Understanding seasonal trends in elemental concentrations, patterns of elemental concentrations over multiple years, and how elemental concentrations shift as a function of genotype and specific environmental conditions are essential for adapting perennial crops to future climates; however, much work remains.

To date, most studies on genotype-specific effects on plant elemental composition have surveyed organs produced above ground including leaves and seeds; however, elements primarily enter the plant through the roots. If there are genotypic or temporal differences in elemental concentrations of the shoot system, this must reflect, at least in part, activities that are happening below ground. One way to study impacts of roots on elemental compositions of the shoot system is to employ the ancient horticultural manipulation of grafting (Mudge et al., 2009; Warschefsky et al., 2016) as an experimental approach. Grafting is commonly used to connect a shoot system (the scion) to the root system of another, often genetically distinct, individual (the rootstock) resulting in an individual that consists of two distinct genotypes connected through a shared vasculature system (Gaut et al., 2019). Through grafting experiments, root system genotype has been shown to influence scion salt tolerance in *Solanum lycopersicum* and *Cucumis sativus* (Huang et al., 2010; Savvas et al., 2011), increase the uptake of many metals in *Citrullus lanatus* during borate stress (Siamak and Paolo, 2019), and improve uptake efficiency of many macronutrients during times of water stress in *S. lycopersicum* (Sánchez-Rodríguez et al., 2014). These data demonstrate that grafting is a powerful system for understanding root system impacts on elemental concentrations in the shoot system.

Grapevine (*Vitis* spp.) is one of the most well-studied and economically important berry crops in the world. Grafting in grapevine became widespread practice in the late 1800s after the North American aphid phylloxera (*Daktulosphaira vitifoliae* Fitch) was introduced into Europe and began decimating European vineyards (Ordish, 1972; Campbell, 2006) destroyed the root systems of European grapevines by sucking sap from the roots and preventing wound healing. To protect European grapevine root systems from phylloxera, several commercial rootstocks were bred using North American *Vitis* species that are naturally resistant to phylloxera infestation (Walker et al., 1991, 1994; Reisch et al., 2012). Additional grapevine rootstock breeding has targeted rootstocks exhibiting acclimation to different soil types (Bavaresco and Lovisolo, 2015; Ferlito et al., 2020), and vine vigor and yield. In grapevines, grafting influences many phenotypes in the shoot system (Cookson and Ollat, 2013; Berdeja et al., 2015; Corso et al., 2016; Zombardo et al., 2020), including shoot elemental composition (Bavaresco et al., 2003; Lecourt et al., 2015; Migicovsky et al., 2019; Harris et al., 2021). The rootstock influence on shoot elemental composition depends on rootstock genotype (Lecourt et al., 2015; Harris et al., 2021), scion genotype (Ibacache et al., 2009; Verdugo-Vásquez et al., 2021), and local environmental conditions (Bavaresco et al., 2003; Lecourt et al., 2015; Migicovsky et al., 2019). Open questions remain regarding how rootstock genotypes modulate differences in elemental composition, how those effects change over developmental, seasonal, and annual cycles, and whether or not those effects correlate with the local environmental conditions.

In this study, we assessed the influence of rootstock genotype, time of year, and local environmental conditions on leaf elemental composition over the course of three years. We used an experimental vineyard in which the grapevine cultivar ‘Chambourcin’ was growing ungrafted (own-rooted) and grafted to three commercial rootstocks. We surveyed concentrations of 20 elements in the leaves at three time points in the season for three consecutive years, and extracted corresponding environmental data from an on-site weather station. We used these data to address the following questions: 1) How does rootstock genotype influence shoot elemental composition? 2) How does elemental composition vary over time and development, and does rootstock genotype influence this variation? 3) What features of the local, above-ground environment explain leaf elemental variation? Does rootstock genotype influence environmental effects on leaf elemental composition?

## Methods

### Study Design

Our study took place in an experimental vineyard at the University of Missouri’s Southwest Research Center in Mount Vernon, Missouri, USA (37.074167 N; 93.879167 W) (Supplemental Figure 1A-B). The French-American hybrid grapevine ‘Chambourcin’ was grown ungrafted (own-rooted) and grafted to three commercial rootstocks: ‘ 1103P’ (*Vitis berlandieri* Planch. x *Vitis rupestris* Scheele) ‘3309C’ (*V. rupestris* x *Vitis riparia* Michx.), and ‘SO4’ (*V. riparia* x *V. berlandieri*) (Supplemental Figure 1C). Each ‘Chambourcin’/rootstock combination was planted in replicates of four vines as part of a fully randomized block (hereafter referred to as ‘cell’ so as not to be confused with larger ‘blocks’ in the vineyard) design. Each cell of four replicated vines was planted twice per row, once in the front half of the row and once in the back half of the row, resulting in eight vines for each rootstock/scion combination per row. Each row in the vineyard was treated with one of three irrigation treatments: full irrigation (100% replacement of evapotranspiration losses), reduced deficit irrigation (RDI; 50% replacement of evapotranspiration losses), and no irrigation (no replacement of evapotranspiration losses). In total, the ‘Chambourcin’ experimental vineyard included 288 vines: four ‘Chambourcin’/rootstock combinations, replicated in cells of four vines per cell, with two cells per row, planted in a total of nine rows. The vineyard is arranged in six spatial blocks containing the full rootstock x irrigation experimental design. Further descriptions of this vineyard can be found in (Maimaitiyiming et al., 2017; Migicovsky et al., 2019; Harris et al., 2021; Swift et al., 2021).

### Sample Collection and Processing

Sampling was conducted at three phenological stages: anthesis (~50% flowers open, mid-late May), veraison (~50% of berries turned from green to red; late July), and immediately prior to harvest (mid-late September). Sampling took place over three consecutive growing seasons from 2017 - 2019. From each grapevine, we collected three leaves from a single shoot: the youngest fully opened leaf (Y), the approximate middle leaf (M), and the oldest leaf (O); hereafter referred to as ‘leaf position’ given that young leaves are always terminal on a shoot, and old leaves are always basal. Leaves were stored in the field in plastic bags and transported back to the lab in coolers, then transferred to individual coin envelopes and dried in an oven at 50°C for 1-3 days. Once dried, leaves were crushed by hand and 75 - 100 mg of leaf tissue was submitted for processing using an established elemental profiling pipeline (Ziegler et al., 2013) at the Donald Danforth Plant Science Center. Samples were acid digested with nitric acid and a spiked internal standard and analyzed using ICP-MS. Measurements were adjusted for sample weight, instrument drift, and internal standards (Ziegler et al., 2013). The elemental profiling pipeline quantifies concentrations of 20 elements: aluminum, arsenic, boron, calcium, cadmium, cobalt, copper, iron, potassium, magnesium, manganese, molybdenum, sodium, nickel, phosphorous, rubidium, sulfur, selenium, strontium, and zinc.

### Data Preprocessing

Missing data precluded fitting equally parameterized models across all elements. Analytical issues prevented the measurements of sulfur and potassium in harvest 2018 samples and calcium and selenium samples in harvest 2019 samples. To overcome this barrier, we opted to strategically impute values missing due to analytical issues using an optimized imputation pipeline. Samples that contained elements with a concentration more than eight standard deviations from the mean value for that element were removed. Remaining data were centered and scaled using a Z-score transformation. The transformed data set was used to train a *K* Nearest Neighbor (*K*NN) model to infer missing values as implemented in the ‘KNNimputer’ function in SciKit-Learn (Pedregosa et al., 2011). The value of *K* was optimized such that imputation errors on known data were minimized. To do this, we removed all samples that contained missing data, and in 50 iterations, we randomly introduced unknown measurements (NAs) to 30 percent of samples for all values of *K* (2 to 100; step size 2). For each model fit, we computed the squared error for each prediction as compared to its true value, and selected the optimal *K* based on the lowest median squared error per *K*. KNN-imputed data were back-transformed into elemental concentrations for linear models and machine learning.

### Model Fitting

#### General Framework

We used a two-phase statistical framework to determine how each factor of the experimental design (rootstock genotype, year, phenological stage, leaf position, and their interactions) influenced shoot elemental composition. First, we fit a random forest to each factor in order to make predictions about factor labels (rootstock genotype, year, phenological stage, leaf position) given the shoot elemental composition (in model notation: `factor ~ elemental composition`). Second, we fit linear mixed models to each element to understand how the experimental design influenced a particular element (in model notation: `element concentration ~ experimental design`). In this framework, we paid special attention to similarities and differences in the interpretation of various model fits. The approach for each of these phases is described below. Unless otherwise noted, all analyses were carried out in R v3.6.1 (R Core Team, 2019).

#### Random Forests

We fit random forest models to determine the predictability of a factor given a complete set of element concentrations using the python package SciKit-Learn (Pedregosa et al., 2011). Here, we focused on the prediction of the rootstock to which ‘Chambourcin’ was grafted and each of the three time components of the study: year, phenological stage, and leaf position, a correlate for leaf age. The back-transformed KNN-imputed data were randomly split into two partitions: train (80%) and test (20%). For each fit, we used random search and 5-fold cross validation within the training set to optimize six model parameters: number of trees, number of features, maximum number of splits, minimum samples to split a node, minimum samples per leaf node, and method to select samples. From each optimization, we selected the best estimator (the model with highest mean accuracy across folds). This optimized model was used to make predictions on the test set to assess model performance using the model accuracy. In addition, we performed 10-fold cross validation on the best estimator using the training set to determine the importance of each element in each model (based on the Gini importance criterion).

#### Linear Mixed Models - Parameterization and Transformation

Some factors of our experimental design, such as irrigation and block, were known to have minimal influence on leaf element concentrations from prior studies (Migicovsky et al., 2019; Harris et al., 2021; Swift et al., 2021). Because of this, we tested model components to determine how best to parameterize our linear models. We fit a linear discriminant analysis (LDA) to the scaled, KNN-imputed elemental concentration matrix using the R package MASS (Ripley, 2002) for each of the following design factors: phenological stage (known to be significant), irrigation (of mixed results in the past), and block and row (to determine the most appropriate descriptor of spatial variation). Model terms were considered for inclusion in the models by assessing the probability that the prediction accuracy was greater than a model making random predictions using the function confusionMatrix from the R package caret (Kuhn, 2013). From this testing, we included irrigation as a non-interacting fixed effect and accounted for potential spatial variation using block as a random effect.

In addition to model parameterization, we explored common response-variable transformations to identify how to best fit linear models under the assumption of normally distributed residuals. We examined raw element concentration, z-score transformations, square root transformations, and log transformation. For each ion, we fit our parameterized model and examined the skew, kurtosis, and results of a Shapiro-Wilk normality test of the residual distribution. Given the sample size of this study, no transformation returned normally distributed residuals. However, the log transformation consistently returned the residual distribution with the lowest skew and the highest *P*-values for the normality test. As such, each element was modeled by its log concentration.

#### Linear Mixed Models - Fitting and Interpretation

Linear mixed models were fit using a repeated measure framework using the lmerTest implementation of linear mixed models (Kuznetsova et al., 2017). In each model, the response was centered and scaled so that models could be compared. Each ion was fit with the following model: `scale(log(element)) ~ irrigation + year*phenology*rootstock*leaf_position + (1|vine_id) + (1|block)`. *P*-values associated with each term were computed using a type-3 ANOVA adjusted using the false discovery rate (Benjamini Hochberg) correction. For terms that were significant in each model, the proportion of variation explained (eta^2^) by term was computed. If significant variation was explained by a main effect, post-hoc comparisons of estimated marginal means were tested using the R package emmeans (Lenth et al., 2018). Post-hoc comparisons were considered significant if their Tukey-corrected p-values (the default method for within model comparisons in emmeans) were below 0.0025 (a Bonferonni alpha correction accounting for 20 elemental models). P-values reported as ‘0’ were below the MacOS system reporting limit of R (2.2e-308). In all other cases, exact p-values were reported.

### Environmental Analysis

Over the course of the experiment, vines experienced natural environmental variation. An on-site weather station (University of Missouri Extension) captured hourly weather data and summarized daily composites for 12 features of the local weather: minimum daily temperature (minT), maximum daily temperature (maxT), average daily temperature (avgT), maximum daily wind speed (maxW), daily precipitation (precip), maximum daily pressure (maxP), minimum daily pressure (minP), maximum daily dew point (maxDP), minimum daily dew point (minDP), average daily dew pressure (avgDP), total solar radiance (rad), and reference short crop evapotranspiration (evap). In addition, we computed three additional environmental features related to daily total changes in each of the following: temperature (dT), pressure (dP) and dew point (dDP). From these 15 environmental measurements, we constructed composite features summarizing environmental conditions that the vines experienced for each of three sample windows: 1) a two-week window prior to sampling, 2) a one-week window prior to sampling, and 3) a three-day window prior to sampling. Each composite was the average condition experienced by the vine except for precipitation and solar radiance which were summed (sumPrecip and sumRad, respectively). In addition to the composite statistics, we included the measurements taken by the weather station from the day of sampling (excluding computed features as the vines would not have experienced the full range by time of sampling). Within a given sampling window, many of the features were highly correlated (e.g., average daily minimum and maximum pressure), therefore environmental features were clustered for each window size and a representative feature was chosen from each cluster. Then, we estimated the relationship of each representative environmental feature from each cluster to each of the 20 elements assayed in this study after adjusting the concentrations for variation from vine ID and leaf position using a linear mixed model. A subsequent linear model was used with fixed effects for rootstock, environmental feature, and the rootstock by environmental feature interaction. The estimated slope of each linear model represented how much a unit change in the environmental feature influenced the scaled elemental concentration. Because such models were computed against scaled elemental concentration, a slope of 1 would indicate that a unit change in environmental feature would expect to change the elemental concentration by one standard deviation. Significant interactions between rootstock and environmental features suggested different rootstock genotypes had different slopes. Such cases were assessed using the emtrends function of the ‘emmeans’ R package. All three sample windows showed highly similar results, so we report only the results from the one-week window and highlight a unique feature of nickel as it correlated with day-of evapotranspiration measurements.

## Results

The experimental design, which included rootstock genotype, year of sampling, time of season, shoot position, and all possible interactions, explained between 37.0% and 88.8% of variation in leaf elemental composition (Figure 1; Supplemental Tables 1 - 4; Supplemental Figures 2-5). Generally, the temporal and developmental components of the study explained more variation than rootstock genotype. Across all elements, year explained an average of 12.9% of elemental variation (Supplemental Figure 3), phenology explained an average of 13.2% (Supplemental Figure 4), and shoot position, a proxy for leaf age, explained 11.4% (Supplemental Figure 5). The strongest overall influence on shoot elemental composition was the interaction between year and phenology, which explained, on average, 14.0% of elemental variation (Supplemental Figure 6). Adjusted marginal means and 95% confidence intervals can be found for each main effect in this study in Supplemental Figures 2-5, and pairwise percent differences across the main effects for each element can be found in Supplemental Tables 1-4.

**Figure 1:**
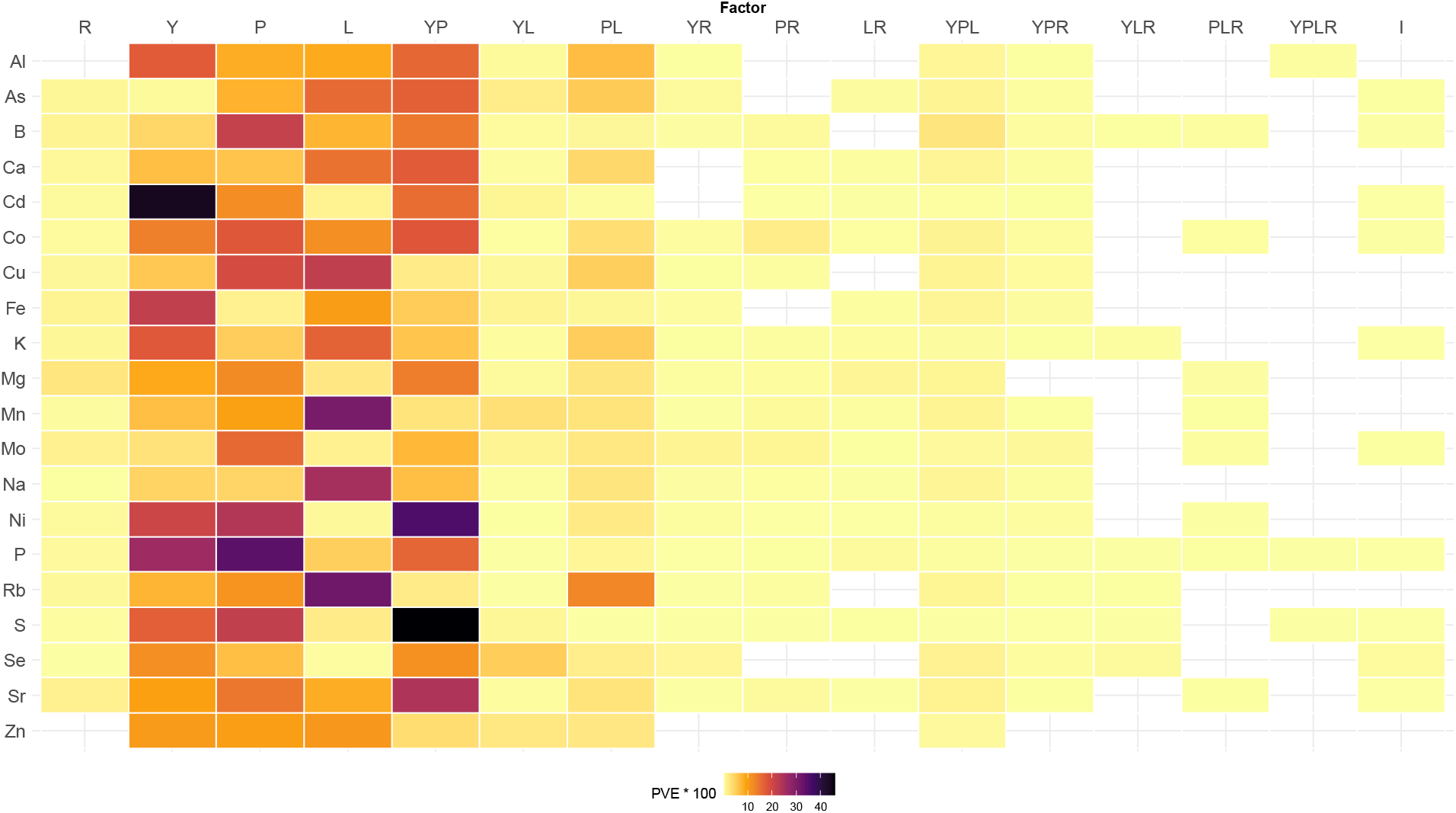
Summary of linear mixed models. A heat map showing the effect size (variation explained) for each term in the linear mixed models (columns) fit to each element (row). Empty cells indicate the term was not significant in the linear model for that element after correcting each associated p-value using the false discovery method. R = Rootstock, Y = Year, P = Phenology, L = Leaf position, I = Irrigation. Interactions are indicated by groups of letters; for example, YP = year × phenology interaction.

### Q1: How does rootstock genotype influence leaf elemental composition?

The rootstock to which ‘Chambourcin’ was grafted was predictable based on elemental composition of the leaf with an accuracy of 72.2% with a random forest (Supplemental Figure 7). Elements most important for this classification were nickel, magnesium, molybdenum, and strontium. Linear mixed models showed that rootstock significantly influenced 18 of the 20 elements surveyed in this study. The only elements that did not vary significantly across rootstocks were aluminum and zinc (Figure 1). Variation explained in the elements significant for rootstock ranged from 0.09% (selenium; p.adj = 0.04) to 2.57% (magnesium; p.adj = 5.52e-47) (Supplemental Table 1). Five elements had more than one percent of variation explained by rootstock, each showing a different general response pattern. Magnesium, which had the strongest effect from rootstock (Figure 2A), had elevated concentrations in both ungrafted and ‘ 1103P’-grafted vines as compared to ‘3309C’- and ‘SO4’-grafted vines. For example, the difference between ungrafted vines and ‘3309C’-grafted vines was 24.51% (p = ‘0’), and the difference between ungrafted vines and ‘SO4’-grafted vines was 23.35% (p = ‘0’) (Supplemental Table 1). The second strongest rootstock effect (1.5%) was observed for strontium (p.adj = 4.20e-42; Figure 2B) which was lower in ‘3309C’-grafted vines relative to ungrafted vines, ‘ 1103P’-grafted vines, and ‘SO4’-grafted vines. For molybdenum (1.46%; p.adj = 7.87e-27; Figure 2C), ‘1103P’- and ‘SO4’-grafted vines had elevated concentrations relative to ungrafted and ‘3309C’-grafted vines. The only element for which the random forest and the linear mixed models differed with respect to elemental importance was nickel. Nickel was the most important element identified in the random forest, but ranked 13th for variation explained by rootstock in the linear mixed model (0.47%; p.adj = 3.04e-33; Figure 2D).

**Figure 2:**
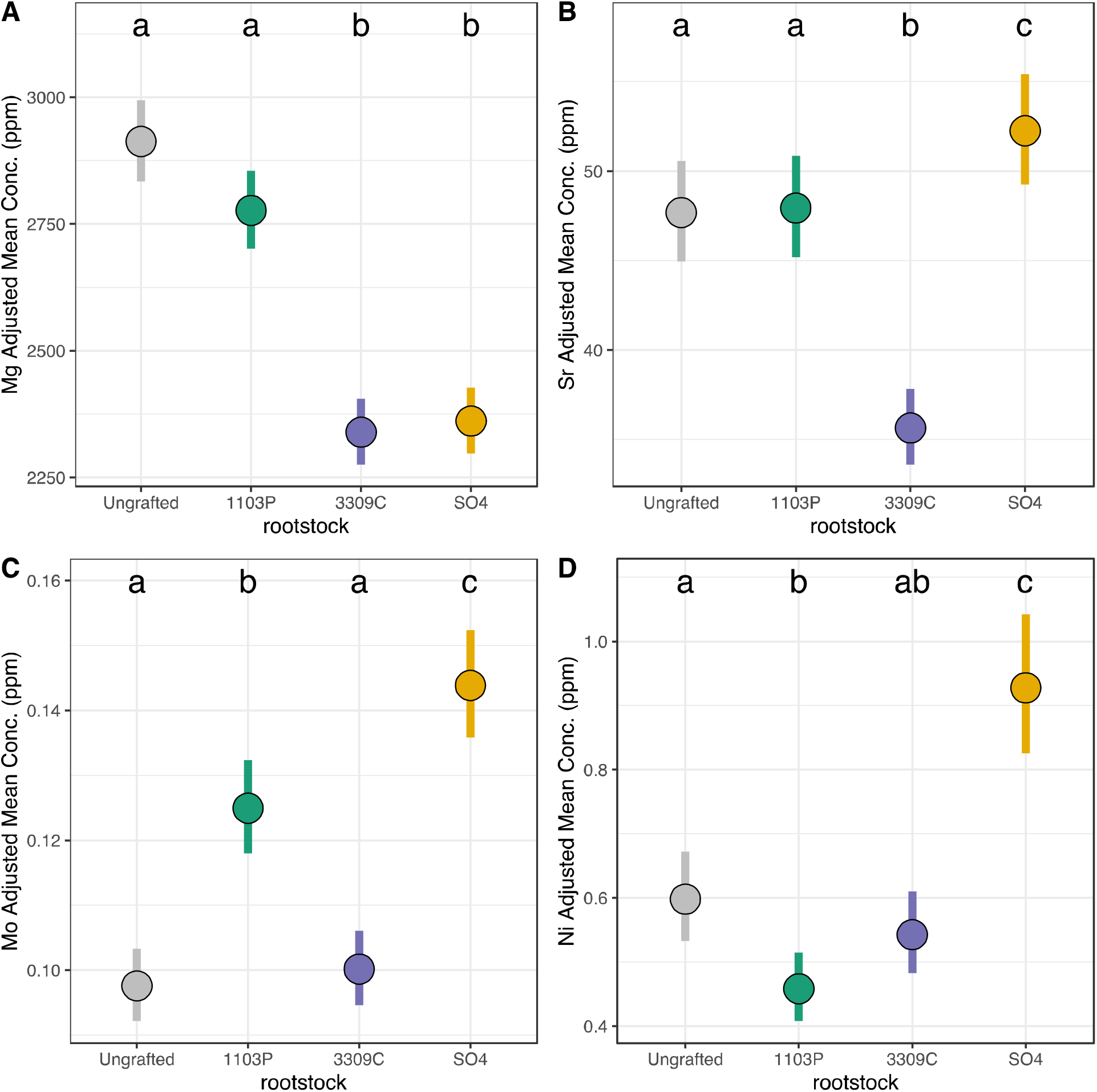
Most variable elements by rootstock genotype. Model-adjusted means and confidence intervals back-transformed into concentrations for elements responding most strongly to the rootstock treatment: **A.** Magnesium (Mg), **B**. Strontium (Sr), **C**. Molybdenum (Mo). Additionally, we show the model adjusted means of the most important elements in rootstock prediction: **D**. Nickel (Ni). Letters above each plotted means indicate the results of a post-hoc comparison of estimated marginal means. P-values were corrected for multiple within-model comparisons using a Tukey adjustment, and significance was assessed after Bonferonni correction for multiple models (alpha = 0.0025).

Individual elements showed diverse response patterns across rootstock genotypes (Supplemental Figure 8). The most common pattern, observed in eight of 20 elements and highlighted in the results of the random forest, was that elemental concentrations in leaves from ungrafted ‘Chambourcin’ vines were more similar to leaves from ‘Chambourcin’ grafted to ‘3309C’ than to leaves of ‘Chambourcin’ grafted to ‘1103P’ or ‘SO4’ (arsenic, cadmium, cobalt, manganese, molybdenum, phosphorus, rubidium, and sulfur) (Supplemental Figure 8A; Supplemental Table 1; Figure 2C, for example).Other detectable patterns included increased concentrations of copper and iron in ‘ 1103P’-grafted ‘Chambourcin’ relative to other root/shoot combinations (Supplemental Figure 8B), decreased concentrations of calcium and strontium in ‘3309C’-grafted vines (Supplemental Figure 8C), and increased concentrations of calcium, molybdenum and nickel in ‘SO4’-grafted vines (Supplemental Figure 8C, Figure 2C, Figure 2D]).

### Q2.1: How does leaf elemental composition vary over time and development?

#### Year

The year from which a leaf was sampled was highly predictable based on elemental composition of the leaves at 97.6% accuracy with a random forest (Figure 3A). Elements most important for the prediction of year were cadmium, nickel, and phosphorus (Figure 3A). Linear mixed models showed that all 20 elements were influenced by the year of sampling (Figure 1; Supplemental Figure 3). Four elements had over 20% of variation explained by year: cadmium (43.6%), phosphorus (26.2%), iron (21.9%), and nickel (20.6%) (Figure 1; Supplemental table 2). For these four elements, the adjusted p-values are below the reporting limit (p.adj = ‘0’). Both the random forest and the linear mixed models converged on identifying cadmium, phosphorus, and nickel as the most significantly affected by the sampling year.

**Figure 3:**
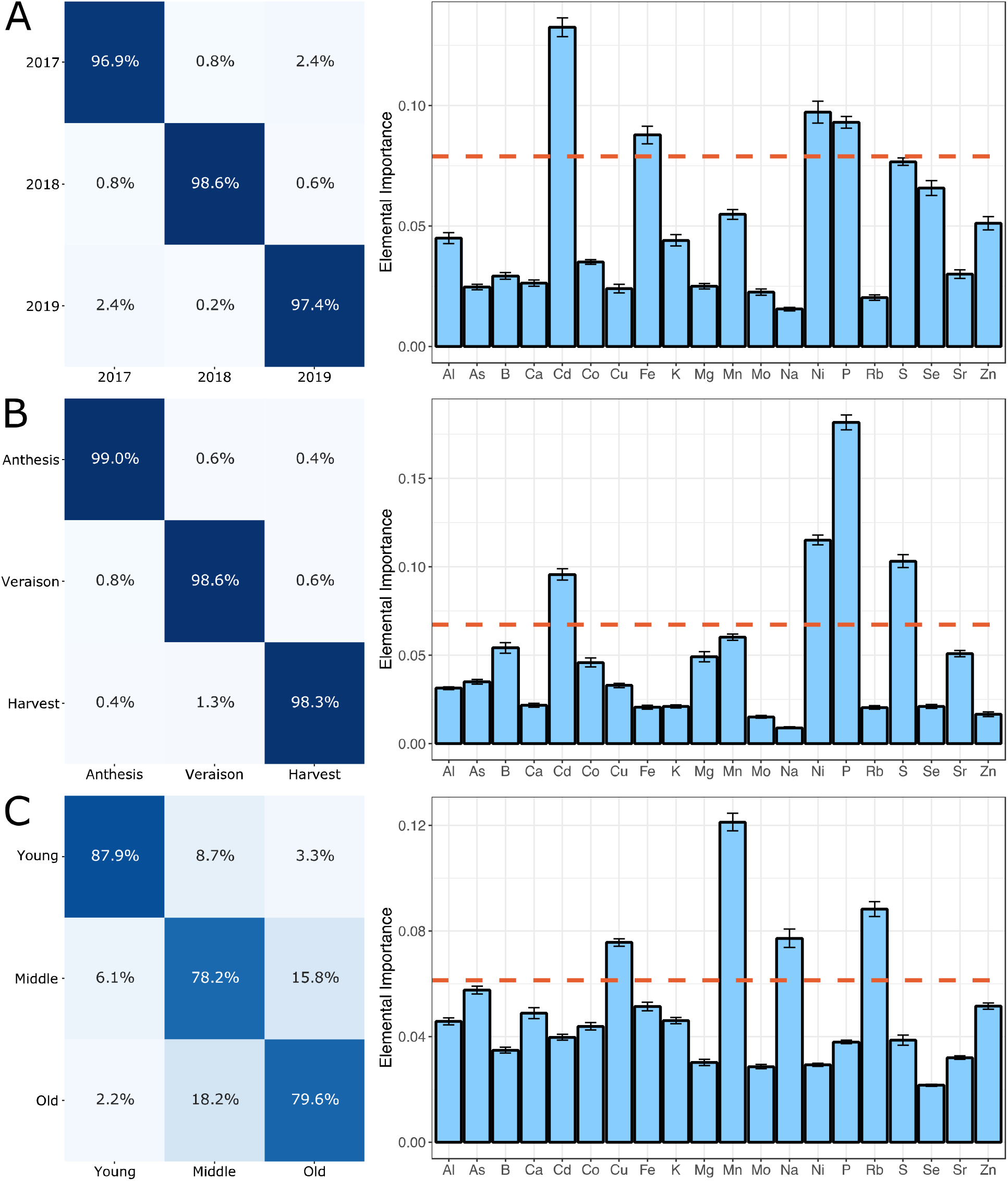
Summary of random forests model to time components. Confusion matrices and estimates of feature importance for the predictions of each time component in our study: **A.** Year, **B.** Phenology), and **C.** Leaf Position. In each case, the confusion matrices are shown for a 20% withheld test set, and the feature importances are derived from 10-fold cross validation of the 80% training set. The orange line represents the 80th percentile for all estimated importances.

Across all three sampling years, the primary patterns that emerged were that elemental concentrations either peaked or were significantly reduced in 2018 (Supplemental Figure 9A; Supplemental Table 2). Ten of 20 elements, including nickel, were at their highest concentration in 2018 and lower concentrations in 2017 and 2019. In comparing 2017 and 2018 adjusted concentrations of nickel, there was a 223.14% increase between the adjusted mean concentrations in 2017 and 2018. Between 2018 and 2019, there is a 52.80% decrease in adjusted nickel concentration. Similarly, eight of 20 elements, including phosphorus and cadmium, were lowest in 2018 and higher in 2017 and 2019. In phosphorus, there was a 44.46% decrease in concentration between 2017 and 2018 and a 116.53% increase between 2018 and 2019. The two elements that did not show a marked change in 2018 were molybdenum and calcium. Molybdenum concentration decreased persistently over the course of the study (10.34% decrease from 2017 to 2018 and an additional 7.87% decrease from 2018 to 2019). Calcium concentrations increased 28.19% from 2017 to 2018, but the comparison between 2018 and 2019 was not significant.

#### Phenology

The phenological stage from which a leaf was collected (anthesis - spring; veraison - midsummer; harvest - fall) was the most predictable factor in this study, with a random forest prediction accuracy of 98.6% (Figure 3B). The most important elements in this model were phosphorus, nickel, and sulfur (Figure 3B). As a single fixed effect in linear mixed models, phenology significantly influenced all 20 elements in the study (Figure 1; Supplemental Table 3, Supplemental Figure 4). Variation explained by phenological stage ranged from 1.42% (iron) to 34.1% (phosphorus). Similar to the elements considered most important in the random forest model, the next highest amounts of variation explained came from nickel and sulfur, 23.7% and 21.9%, respectively (Supplemental Figure 4).

There were several general response patterns of concentration variation across phenological time points (Supplemental Figure 9B; Supplemental Table 3). The first common pattern (seven of 20 elements; including nickel) showed the concentration of the element was highest at veraison, and lower in anthesis and harvest. For example, nickel concentration increased 21.85% between antheses and veraison and then decreased 70.51% between veraison and harvest. The second common pattern (seven of 20 elements; including phosphorus and sulfur) showed persistent decreases in concentration over the course of the season. For example, phosphorus concentrations decreased 49.33% from anthesis to veraison and 17.35% from veraison to harvest. Similarly, sulfur concentrations decreased 10.62% from anthesis to veraison and 83.95% from veraison to harvest. Finally, the remaining six elements fell into two general response patterns: a decrease between anthesis and veraison and then no significant change between veraison and harvest (four elements) and a decreased concentration at veraison relative to both anthesis and harvest (two elements).

#### Leaf Position

The leaf position along a shoot was predictable by random forest with an accuracy of 81.9% (Figure 3C). The most important elements in the prediction of leaf position were manganese, rubidium, sodium, and copper (Figure 3C). Linear mixed models showed that all elements were significantly influenced by leaf position with variation explained ranging between 0.31% (selenium) and 31.85% (rubidium) (Figure 1; Supplemental Table 4; Supplemental Figure 5). Linear mixed models confirmed the elemental importance of the random forest by identifying the same top four elements as having significant variation explained by leaf position: rubidium, manganese (30.76%), sodium (25.38%), and copper (22.09%) (Supplemental Figure 5). For all four of these elements, the adjusted p-value associated with leaf position was below the reporting threshold (p.adj = ‘0’).

Elemental responses to leaf position grouped into four general response patterns (Supplemental Figure 9C; Supplemental Table 4). The most common response pattern showed decreased concentrations in the middle leaf position relative to younger and older leaves (seven out of 20 elements; including sodium). Elements grouped into this general pattern differed where the highest concentration was found. For example, three elements (aluminum, cadmium, and iron) showed the highest concentration in older leaves, while boron, sodium, phosphorus, and zinc had the highest concentration in younger leaves. In sodium, there was a 61.07% decrease in concentration from younger to middle leaves and 24.13% increase from middle to older leaves. The second most common response pattern showed persistent increases in concentration with leaf age (five elements). In manganese, there was a 29.37% increase in concentration between young and middle leaves and a 86.66% increase in concentration between middle and old leaves. The third most common response pattern showed a persistent decrease in elemental concentration with leaf age (four elements). For example, in rubidium, there was a 52.28% decrease between young and middle leaves and a 21.51% decrease between middle and old leaves. Similarly, there was a 39.76% decrease in copper concentration between young and middle leaves and a 20.26% decrease between middle and old leaves. All other elements were not significantly different between young and middle leaves and saw increases in concentrations between middle and older leaves (four elements).

### Q2.2: Does rootstock genotype influence temporal variation in elemental composition?

The rootstock by year interaction significantly influenced the concentrations of 17 out of 20 elements. Variation explained by the interaction of rootstock and year ranged from 0.06% (phosphorous, p.adj = 2.05e-04) to 1.17% (molybdenum, p.adj = 8.20e-25). In grafted vines, molybdenum concentrations decreased across the three years of this study (Figure 4A). However, in ungrafted vines, the concentration of molybdenum increased between 2018 and 2019. Following molybdenum, the element with the second most explained variation by the interaction of rootstock and year was selenium (0.88%; p.adj = 1.27e-16; Figure 4A). Selenium in all rootstock genotypes peaked in concentration in 2018, but the rank order changed such that ungrafted vines and vines grafted to ‘3309C’ were highest in concentration in 2017, but lowest in concentration in 2018. In 2019, no rootstock genotypes were significantly different from one another. The interaction of rootstock with year was not significant for calcium, cadmium, or zinc.

**Figure 4:**
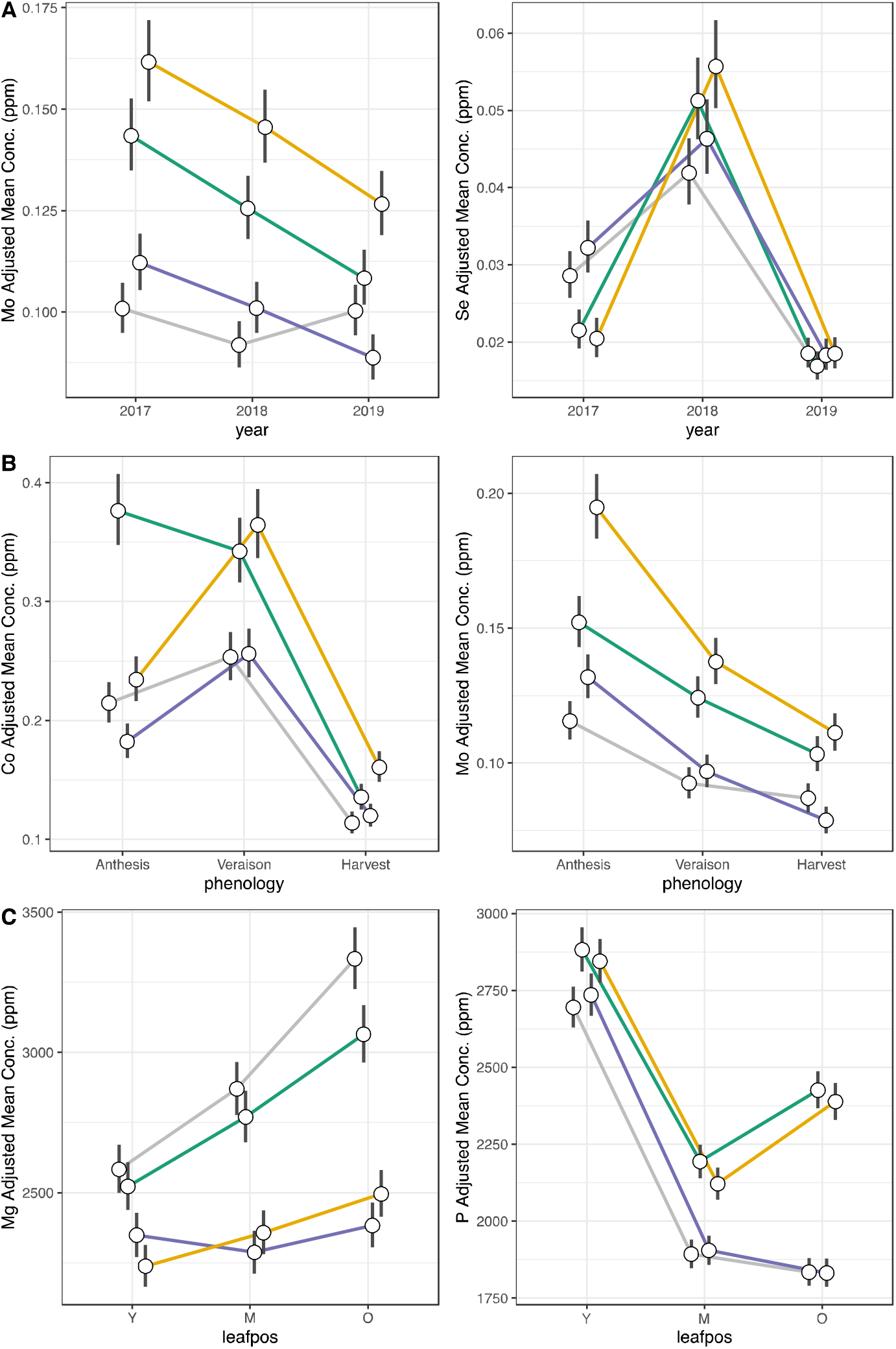
Rootstock by time interactions Model-adjusted means and 95% confidence intervals back-transformed into concentrations for elements responding most strongly to the rootstock by time interactions of our study: **A.** Year (cobalt and molybdenum), **B**. Phenology (molybdenum and selenium), **C**. Leaf position (magnesium and phosphorous). Gray lines represent ungrafted vines, teal lines represent ‘ 1103P’-grafted vines, purple lines represent ‘3309C’-grafted vines, and yellow lines represent ‘SO4’-grafted vines.

The interaction between rootstock and phenological stage was significant for 15 out of 20 elements. For elements significantly influenced by this interaction, the amount of variation explained ranged from 0.04% (nickel; p.adj = 5.54e-03) to 1.81% (cobalt; padj = 1.40e-83). At anthesis, cobalt concentrations in vines grafted to ‘ 1103P’ were significantly higher than all other rootstock genotypes (Figure 4B). At veraison, both ‘ 1103P’- and ‘SO4’-grafted vines had significantly higher concentrations compared to ungrafted and ‘3309C’-grafted vines. By harvest, cobalt concentrations were considerably decreased in all rootstock genotypes, but both ‘1103P’- and ‘SO4’-grafted vines were enriched compared to the other genotypes. Following cobalt, the rootstock by phenology interaction most strongly influenced molybdenum concentrations (1.06%; p.adj = 3.09e-22; Figure 4B). All grafted vines showed decreases in molybdenum as the season progressed; however, ungrafted vines did not show a decrease between veraison and harvest. Rootstock interaction with phenology was not significant for aluminum, arsenic, iron, strontium, and zinc.

The interaction between rootstock and leaf position was significant for 14 out of 20 elements. Variation explained by the interaction of rootstock and leaf position ranged from 0.07% (nickel; p.adj = 1.23e-04) to 0.96% (magnesium; p.adj = 5.97e-24). Magnesium tended to accumulate in leaves as they aged (Figure 4C). However, ‘Chambourcin’ grafted to ‘3309C’ showed no change in concentration between young leaves and middle leaves. Following magnesium, phosphorus (0.51%; p.adj = 1.62e-45; Figure 4C) had the second highest amount of variation that could be explained by the interaction of rootstock and leaf position. All ‘Chambourcin’ vines showed sharp decreases in phosphorus concentrations between young and middle leaves. Between middle and older leaves, ungrafted vines and ‘3309C’-grafted vines experienced a small decrease in phosphorus concentration; however, vines grafted to ‘ 1103P’ and ‘SO4’ showed marked increases in phosphorus concentrations. The interaction of rootstock and leaf position was not significant for aluminum, boron, copper, rubidium, selenium, and zinc.

### Q3: Environmental Variation

Each of the 20 elements measured were significantly influenced by at least one of the environmental features used in the linear models (Figure 5A-B). Moreover, each environmental feature had a subtly different influence on elemental composition overall (Figure 5B). Generally, the strongest influence came from the cumulative amount of precipitation (sumPrecip), average daily maximum pressure (maxP), and the average daily change in temperature (dT). For example, phosphorus had 64.6 % of its variation explained by the cumulative amount of precipitation (Figure 5C), 56.3% of the variation in rubidium was explained by the average daily maximum pressure (Figure 5C), 31.8% of the variation in manganese was explained by the average daily change in temperature (Figure 5C), and 24.8% of the variation in cobalt was explained by the average daily change in dew point (dDP). Average daily estimated reference crop evapotranspiration (Evap) explained significant variation in more than any other environmental feature (16 out of 20) with the highest proportion being in strontium, rubidium, potassium, and aluminum.

**Figure 5:**
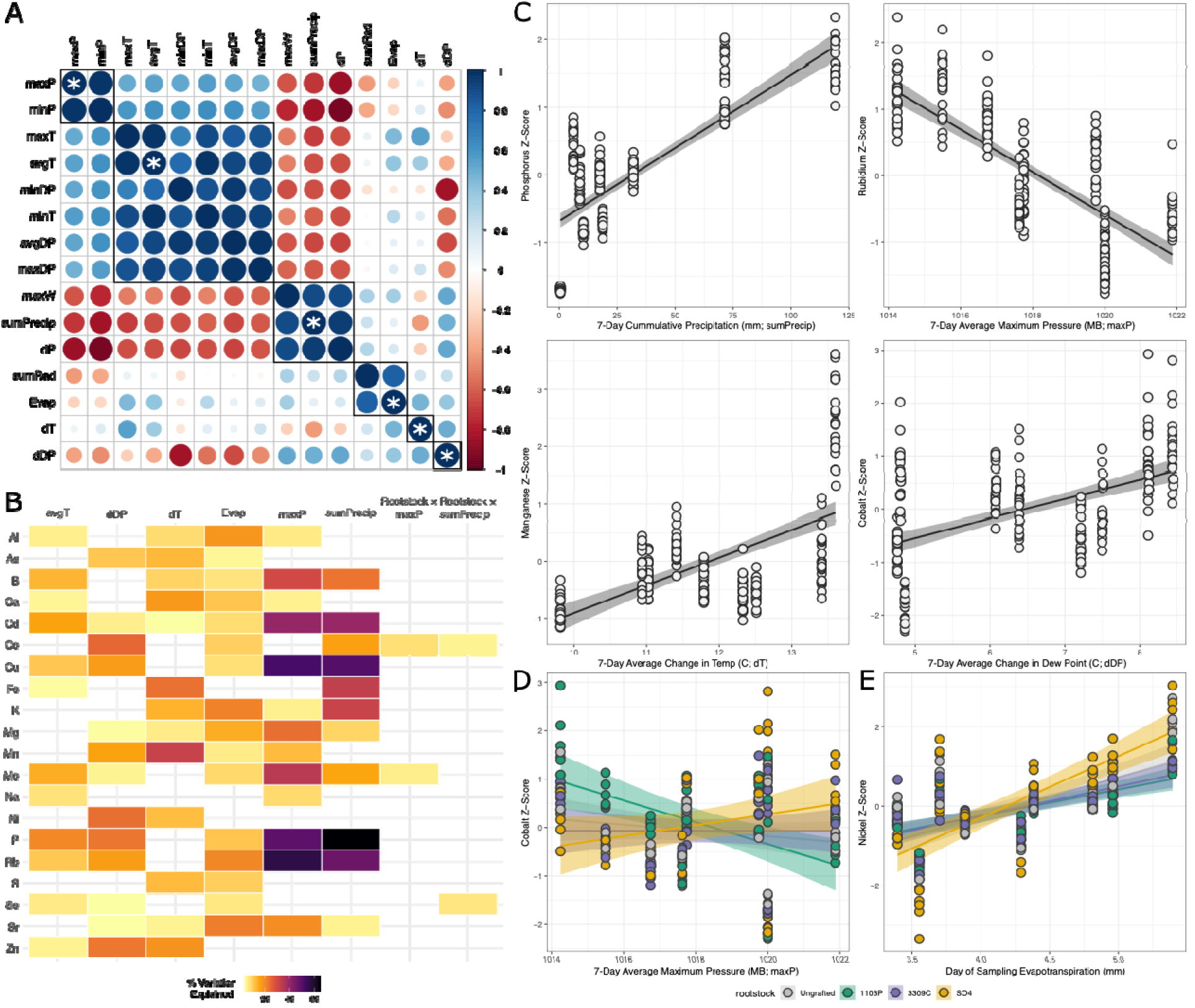
Elemental composition over environmental variation. **A.** Correlation matrix (correlogram) of each seven-day composite environmental features derived from the on-site weather station. The strength of the correlation is indicated by the size and color intensity of the circle. Black boxes indicate the results of hierarchically clustering the correlation matrix and cutting at the tree such that six modules would be identified. The environmental feature selected for linear modeling is shown with a white asterisk. **B.** Summary of linear models fit to each element with each seven-day composite environmental features and rootstock genotype x environmental features included in the model. In total, this figure summarizes six linear models fit to REML adjusted concentration, removing variation from vine id and leaf position. Model terms and cells missing from the figure were not significant after adjusting for multiple tests (fdr). The intensity of each cell in the figure shows the proportion of variation explained by that model term (column) for a particular element (row). **C.** Elements significantly correlated with environmental variation. Elements were selected for visualization by sorting on variation explained and making selections such that each element and each environmental feature were only shown once. X-axes are labeled with full descriptions of the environmental feature and the shortened names that connect them to panels A-B. **D.** Cobalt, the element with the most variation explained by the interaction of rootstock and seven-day composite environmental features. Here is it shown against seven-day average maximum pressure. Points are colored by rootstocks, and the lines are fit by rootstock. The X-axis is labeled with full descriptions of the environmental feature and the shortened name that connects it to panels A-B. **E**. Nickel, an example element significant for day-of environmental features. Here it is shown against day-of reference short crop evapotranspiration. Points are colored by rootstocks, and the lines are fit by rootstock. Note: this panel is not directly connected to Panels A-B.

In four elements, there was evidence of a rootstock-mediated effect of the environment on elemental composition (Figure 5D). Cobalt showed the strongest mediated effects with 7.5% of its variation being explained by the interaction of rootstock and average daily max pressure. Vines grafted to ‘SO4’ had a significantly larger slope (0.06) than vines grafted to ‘1103P’ (−0.14; p = 0.0002). Selenium showed a similarly large effect (6.6%) with cumulative precipitation. Ungrafted vines had a significantly higher slope (0.002) then vines grafted to ‘1103P’ (−0.001; p = 0.004) and vines grafted to ‘SO4’ (−0.001, p = 0.006). Molybdenum was significant for the interaction of rootstock and the change in temperature on the day of sampling (5.1%). Ungrafted vines (slope = −0.04) had a significantly higher slope than vines grafted to ‘3309C’ (slope = −0.16, p = 0.007) and vines grafted to ‘SO4’ (slope = −0.18, p = 0.001). Nickel showed significant variation (4.9%) explained by the interaction of rootstock and day-of-sampling estimated reference crop evapotranspiration. All rootstocks showed significant positive slopes between nickel concentration and evapotranspiration, but vines grafted to ‘SO4’ showed a significantly stronger relationship. Post-hoc comparisons showed that the relationship between nickel and evapotranspiration was higher in ‘SO4’ (slope = 1.02) than vines grafted to ‘1103P’ (0.44, p = 0.0005) and ‘3309C’ (slope = 0.48, p = 0.002).

## Discussion

In this study, we used grafted grapevines to understand the influence of rootstock genotype on leaf elemental composition over multiple years, phenological stages, and leaf ages. Expanding on previous work, we showed that rootstock genotype adds an additional layer of complexity on top of inter-annual variation and expected seasonal and leaf age progressions. Moreover, we found that leaf elemental composition varies significantly with many aspects of the above-ground environment including features related to pressure, temperature, and precipitation. Finally, we show that in some cases, elemental composition varies through the interaction of rootstock genotype and above-ground environment suggesting that grafted plants are sensing changing environmental conditions and coordinating a response to drive changes in element uptake.

### Leaf elemental composition varies as a function of rootstock genotype

Previously, we identified an effect of rootstock genotype on shoot elemental composition in ‘Chambourcin’ using leaves collected near anthesis (Migicovsky et al., 2019), and that this effect was variable across the season (Harris et al., 2021). Here, we used an updated modeling framework to account for variation between vines and two additional years of data to show that the rootstock genotype most heavily influenced magnesium, strontium, and molybdenum concentrations. Moreover, we showed again that nickel was the most important element for predicting rootstock genotype due to a large difference between the concentration of nickel in the leaves of vines grafted to ‘SO4’ as compared to the other graft treatments in this study. The influence of rootstock genotype on shoot elemental composition is not unique to ‘Chambourcin’ and has been shown in other grapevine cultivars (Bavaresco et al., 2003; Lecourt et al., 2015); however, direct comparisons have been difficult due to differences in rootstock and scion genotypes used in previous studies.

Rootstock-mediated differences in leaf elemental composition of a common scion such as ‘Chambourcin’ point to the existence of specific features within the rootstock genotype that contribute to differences in ion uptake. A study of ‘Cabernet Sauvignon’ grafted to 13 commercial rootstock cultivars in a common vineyard showed each of the three commonly used North American species for rootstock breeding imparted a unique signature onto shoot elemental composition (Gautier et al., 2020). For example, scions grafted to rootstocks with *V. riparia* in its pedigree conferred more sulfur and less phosphorus, magnesium, boron, and aluminum to shoot petioles. Interestingly, our study, while not designed to address this question, provides evidence that some of these effects are consistent across study sites. We found that rootstocks with *V. riparia* in their pedigrees conferred less magnesium to ‘Chambourcin’ leaves, relative to ungrafted ‘Chambourcin’ leaves or those grafted to rootstocks that do not have *V. riparia* in their pedigrees. Rootstock pedigree could explain the most common rootstock effect that we observed in this study, where vines grafted to ‘ 1103P’ tended to be more similar to vines grafted to ‘SO4’. Both ‘ 1103P’ and ‘SO4’ share *V. berlandieri* as a parent, but *V. berlandieri* is not present in the pedigree of ‘3309C’ or ‘Chambourcin’ (for ungrafted vines). In the model system *Arabidopsis*, genetic mutants used as rootstocks have been shown to influence leaf elemental composition (Baxter et al., 2008, 2009; Tian et al., 2010), and our findings support the idea that there is likely a genetic component within grapevine rootstocks that determines elemental composition in grafted scions. However, future studies will be needed to determine which genes and which species in the pedigree can drive these changes.

### Leaf elemental composition varies over temporal scales

In the traditional model of elemental accumulation for perennials, elements like nitrogen, phosphorus, and potassium are shuttled out of senescing leaves and stored in perennial structures (Sánchez□Alonso and Lachica, 1987a, 1987b; Roca-Pérez et al., 2006). Some elements, such as calcium, manganese, and magnesium, do not follow this model as they have low rates of phloem loading and mobility (Sánchez□Alonso and Lachica, 1987a, 1987b; Maier and Chvyl, 2002; Roca-Pérez et al., 2006; Navarro et al., 2008; Holzapfel, 2019). Our findings support these previously established patterns in grafted grapevines. Moreover, we add to this that other elements followed similar trajectories such as cobalt and strontium which increased in concentration with leaf age.

While patterns of elemental composition are readily apparent across leaf age, they are not all consistent across seasonal progression. For example, both phosphorus and potassium showed decreases in concentration across leaf ages and over the course of the season, but other elements, especially those that tended to increase in concentration with leaf age, showed higher concentrations at veraison. Veraison is a middle-season developmental time point in grapevine where berry development switches from berry growth, characterized by cell division, to berry ripening, characterized by accumulation of sugar, decrease in acid concentrations, and, in red grapes, change in color (Coombe and McCARTHY, 2000). Previously, we showed that the leaves rapidly shift from early season to late season transcriptomes (Harris et al., 2021). However, the current results suggest that there is a unique signature of veraison in ‘Chambourcin’ characterized by elevated leaf concentrations of aluminum, calcium, cobalt, magnesium, nickel, sulfur, and strontium. This observation raises a host of questions including: Do these elements serve unique functions during veraison? How are supposedly phloem-immoble elements like calcium and magnesium excluded from the leaves between veraison and harvest? Is this consistent across different locations/rootstocks/scions? This study highlights the dynamic nature of elemental concentrations within the plant organismal system.

### Above-ground environmental factors interact with rootstock genotype and temporal components to shape leaf elemental composition

Our study builds on previous studies by examining leaf elemental composition with paired, realtime features of the above-ground environment. Most studies that have examined the influence of the environment on elemental composition have fallen into one of three categories: studies that analyzed large-scale climatic features (such as annual rainfall) (Sardans et al., 2015; Luo et al., 2016), studies that compared sites without focus on environmental differences (Asaro et al., 2016; Shakoor et al., 2016), and studies that manipulated the plant’s local environment (e.g., irrigation or salt treatments) (Urbina et al., 2015; Temme et al., 2019; Chavarria et al., 2020; Shelden et al., 2020). Often, effects from these studies tended to be variable with large influence from phylogeny (Neugebauer et al., 2018). While causal links between measurements captured in the present study and changes in elemental composition are beyond the scope of this study, there are clear pathways for causal relationships to be determined. For example, shifts in the environment can lead to changes in soil chemistry (Chadwick and Chorover, 2001) which manifest in the forms of changes in acid/base dynamics, redox potential, and elemental exchange capacity. Longer term experiments with comprehensive, coordinated collection of phenotypic data, above-ground environmental data, below-ground environmental data, and soil composition can help elucidate causal relationships by decoupling correlations in environmental features, removing collinearity with programmed seasonal development, and determining whether such responses are extensible across species or different soil chemistries.

Interactions observed here between rootstock genotype and environmental features suggest that root systems mediate plant environmental responses. We observed significant interactions of rootstock genotype and environmental features for four elements: cobalt, molybdenum, selenium, and nickel. Of particular note, we have previously shown that nickel concentrations in ‘Chambourcin’ leaves display a very strong signature of rootstock genotype. Here we showed that nickel was significantly correlated with day-of measurements of reference short crop evapotranspiration. When accounting for rootstock genotype, vines grafted to ‘SO4’ had a significantly stronger relationship between nickel concentration and evapotranspiration than other rootstocks. Reference short crop evapotranspiration (ET_0_) is a predicted value for how much water a small grass crop would transpire in the given environment (Allen and Food and Agriculture Organization of the United Nations, 1998). ET_0_ is a composite statistic merging information on solar radiance, wind, pressure, and the energetics of heat and vapor transfer in the atmosphere. ET_0_ can be specified to a particular crop (ET_C_) via linear transformation with a crop coefficient, which has been estimated in grapevine to be between 0.5 and 1.08 (Pruitt, 1977; Williams et al., 2003; Campos et al., 2010; Marras et al., 2016; Munitz et al., 2019). While ET_0_ can be useful to estimate water usage, it does not necessarily reflect in-the-field water usage. For example, we previously showed that leaf transpiration and stomatal conductance were not different in ‘SO4’-grafted vines when compared to ungrafted or ‘ 1103P’-grafted vines (Harris et al., 2021). Our results suggest that either: 1) nickel is being accumulated in ‘SO4’-grafted vines because it is required in a mechanism to maintain water balance or 2) nickel is being accumulated in response to something correlated with ET_0_ that was not explicitly measured in this study. The role of nickel in maintaining water balance is not supported by previous literature except in the case of *Thlaspi* spp. where a hyperaccumulating species showed decreased water loss in toxic metal conditions (Kramer et al., 1997), but nickel hyperaccumulation did not enhance survival to severe drought in later studies (Whiting et al., 2003).

## Conclusions

Grafted grapevines offer a powerful experimental tool for understanding how leaf elemental composition reflects coordinated activities of genetically distinct root and shoot systems, and how those interactions shift over years, across phenological stages within a season, and over developmentally distinct positions in the plant. Results presented here highlight how the complex, dynamic processes by which rootstock and scion interact in element uptake and transport throughout the vine are mediated by local environmental conditions. This work further underscores the importance of multi-year studies, incorporating paired phenotypic and environmental data, to understand genetic and environmental underpinnings of perennial plant form and function.

## Supplemental Tables

Supplemental Table 1: Summary of Rootstock Main Effect.

For each element measured in this study, we report the variation explained in the linear mixed model, model-adjusted means for each rootstock genotype, percent difference for each pairwise rootstock comparison, and the emmeans-derived p-value for each pairwise estimate of mean difference between rootstocks.

Supplemental Table 2: Summary of Year Main Effect.

For each element measured in this study, we report the variation explained in the linear mixed model, model-adjusted means for each year, percent difference for each pairwise year comparison, and the emmeans-derived p-value for each pairwise estimate of mean difference between year.s

Supplemental Table 3: Summary of Phenology Main Effect.

For each element measured in this study, we report the variation explained in the linear mixed model, model-adjusted means for each phenological stage, percent difference for each pairwise phenological comparison, and the emmeans-derived p-value for each pairwise estimate of mean difference between phenological stages.

Supplemental Table 4: Summary of Leaf Position Main Effect.

For each element measured in this study, we report the variation explained in the linear mixed model, model-adjusted means for each leaf position/age, percent difference for each pairwise leaf position comparison, and the emmeans-derived p-value for each pairwise estimate of mean difference between leaf positions.

## Supplemental Figure Legends

Supplemental Figure 1: Study design.

**A**. Vineyard Map. The experimental vineyard features 288 vines either ungrafted (own-rooted) or grafted to one of three rootstocks (‘ 1103P’, ‘3309C’, or ‘SO4’). Each row in the vineyard is treated to one of three irrigation treatments (None, 50% reduced deficit (Partial) or Full). **B**. Explanation of sampling. Each cell in the vineyard contained four replicated ‘Chambourcin’/rootstock combinations. From each cell, each replicate was sampled for elemental profiling. From each replicate, we sampled three leaf positions: young, middle, and old. This was done for three timepoints in a single season (anthesis, veraison, and harvest) for three years (2017 - 2019). **C**. Description of rootstock design. Our study features three rootstocks: 1103P, 3309C, and ‘SO4’. Each of these is a cross between (two of) either *V. berlandieri, V. riparia*, or *V. rupestris*. Common parents between each rootstock are shown. This figure is partially reproduced from (Migicovsky et al., 2019) which is available under a Creative Commons license (CC BY 4.0).

Supplemental Figure 2: All elements across rootstock genotypes.

All 20 elements are shown with means and 95% confidence intervals across the four ‘Chambourcin’/rootstock combinations in this experiment. Means were derived from a linear mixed model and adjusted for random variation in vine id and block. Means and confidence intervals are converted from their transformed values back into concentrations. Letters above each plotted means indicate the results of a post-hoc comparison of estimated marginal means. P-values were corrected for multiple within-model comparisons using a Tukey adjustment, and significance was assessed after Bonferonni correction for multiple models (alpha = 0.0025).

Supplemental Figure 3: All elements across years.

All 20 elements are shown with means and 95% confidence intervals across the three years in this experiment. Means were derived from a linear mixed model and adjusted for random variation in vine id and block. Means and confidence intervals are converted from their transformed values back into concentrations. Letters above each plotted means indicate the results of a post-hoc comparison of estimated marginal means. P-values were corrected for multiple within-model comparisons using a Tukey adjustment, and significance was assessed after Bonferonni correction for multiple models (alpha = 0.0025).

Supplemental Figure 4: All elements across phenological stages.

All 20 elements are shown with means and 95% confidence intervals across the three phenology stages in this experiment. Means were derived from a linear mixed model and adjusted for random variation in vine id and block. Means and standard deviation are converted from their transformed values back into concentrations. Letters above each plotted means indicate the results of a post-hoc comparison of estimated marginal means. P-values were corrected for multiple within-model comparisons using a Tukey adjustment, and significance was assessed after Bonferonni correction for multiple models (alpha = 0.0025).

Supplemental Figure 5: All elements across leaf position.

All 20 elements are shown with means and 95% confidence intervals across the leaf positions/ leaf ages in this experiment. Means were derived from a linear mixed model and adjusted for random variation in vine id and block. Means and standard deviation are converted from their transformed values back into concentrations. Letters above each plotted means indicate the results of a post-hoc comparison of estimated marginal means. P-values were corrected for multiple within-model comparisons using a Tukey adjustment, and significance was assessed after Bonferonni correction for multiple models (alpha = 0.0025).

Supplemental Figure 6: All elements over the year by phenology interaction

All 20 elements are shown with means and 95% confidence intervals across the interaction of year and phenology. Means were derived from a linear mixed model and adjusted for random variation in vine id and block. Means and standard deviation are converted from their transformed values back into concentrations.

Supplemental Figure 7: Summary of the random forest for rootstock genotype

Confusion matrix and estimates of feature importance for the predictions of rootstock genotype. The confusion matrix summarizes model performance on a 20% withheld test set, and the feature importances are derived from 10-fold cross validation of the 80% training set.

Supplemental Figure 8: Common patterns of elemental composition by rootstock

This figure showcases a subset of the adjusted means plots from Supplemental Figure 3 with an emphasis on common patterns observed across rootstock genotypes. **A.** The most common pattern whereby 8/20 elements show concentrations in leaves grafted ‘ 1103P’ or ‘SO4’ more similar to those grafted to ‘3309C’ and ungrafted. This pattern is showcased in arsenic (As) and rubidium (Rb). **B.** The second most common pattern where some elements show increased concentration in leaves grafted 1103P as compared to other ‘Chambourcin’/rootstock combinations. This pattern is shown for copper (Cu) and iron (Fe). **C.** The third most common pattern in element composition by rootstock where some elements show decreased concentrations in leaves of vines grafted to 3309C. This pattern is shown for calcium (Ca) and strontium (Sr).

Supplemental Figure 9: Common patterns of elemental composition over time.

This figure showcases a subset of the adjusted means plots from Supplemental Figures 2 - 4 with an emphasis on common patterns observed across the time components of our study. **A.** The most common responses across the years in this study. Exemplar elements show elevated concentrations in 2018 (nickel), decreased concentrations in 2018 (cadmium), a persistent drop in concentration over the study (molybdenum), and concentrations which were significantly lower in 2017 (calcium). **B.** The most common patterns across the phenological stages of this study. Exemplar elements show increased concentrations at veraison (nickel), a persistent decrease over the season (phosphorus), increased concentrations only at anthesis (boron), and a decreased concentration at veraison (manganese). **C.** The most common patterns across leaf position in this study. Exemplar elements show decreased concentration in middle position leaves (sodium), a persistent increase in concentration with leaf age (manganese), a persistent decrease in concentration with age (rubidium), and elevated concentration only in older leaves (calcium).

## Author Contributions

This study was conceptualized by ZHN, LLK, MTK, and AJM. All authors contributed to sample collection, sample processing, data curation or formal data analysis. ZNH and JEP contributed to data visualization. The original draft was written by ZNH, JEP, and AJM. All authors contributed to reviewing and editing the manuscript. All authors approve of the submitted version.

## Acknowledgments

This work was supported by the National Science Foundation’s Plant Genome Research Research Program 1546869. We thank the members of the Kovacs Lab at Missouri State University, the Kwasniewski Lab at Missouri University, and the Miller Lab at Saint Louis University for helping to sample for the study. We thank members of the Miller Lab and the Baxter Lab at the Donald Danforth Plant Science Center for sample processing. We additionally thank members of the Miller Lab and Ivan Baxter for thorough editing and review of this manuscript.

## Data and Code Availability

Data and archived code are publicly available on Figshare (Harris, 2022). In addition, all notebooks used for data analysis are available on GitHub (https://github.com/PGRP1546869/mt_vernon_1719_elementalComposition).

